# REV7/FANCV Binds to CHAMP1 and Promotes Homologous Recombination Repair

**DOI:** 10.1101/2021.10.04.463067

**Authors:** Feng Li, Prabha Sarangi, Hanrong Feng, Lisa Moreau, Huy Nguyen, Connor Clairmont, Alan D. D’Andrea

## Abstract

A critical determinant of DNA repair pathway choice is the HORMA protein REV7, a small abundant adaptor which binds to various DNA repair proteins through its C-terminal seatbelt domain. The REV7 seatbelt binds to the REV3 polymerase to form the Polymerase ζ complex, a positive regulator of translesion synthesis (TLS) repair. Alternatively, the REV7 seatbelt binds to SHLD3 in the Shieldin complex, a positive regulator of NHEJ repair. Recent studies have identified another novel REV7 seatbelt-binding protein, CHAMP1 (Chromosome Alignment-Maintaining Phosphoprotein, though its role in DNA repair is unknown. Here, we show that the REV7-CHAMP1 complex promotes homologous recombination (HR) repair by sequestering REV7 from the Shieldin complex. CHAMP1 competes directly with the SHLD3 subunit of the Shieldin complex for a limited pool of C-REV7, thereby inhibiting the REV7-mediated recruitment of the SHLD2 and SHLD1 effector subunits to DNA double strand breaks. CHAMP1 thereby channels DNA repair away from error-prone NHEJ and towards the competing error-free HR pathway. Similarly, CHAMP1 competes with the REV3 component of the POLζ complex, thereby reducing the level of mutagenic TLS repair. CHAMP1 interacts with POGZ in a heterochromatin complex further promoting HR repair. Importantly, in human tumors, CHAMP1 overexpression promotes HR, confers PARP inhibitor resistance, and correlates with poor prognosis. Thus, by binding to either REV3, SHLD3, or CHAMP1 through its seatbelt, the REV7 protein can promote either TLS repair, NHEJ repair, or HR repair respectively.

## INTRODUCTION

REV7 (also known as MAD2L2, MAD2B, or FANCV), is a highly-conserved member of the HORMA family of proteins, named for its three founding members: <u>HOp1</u>, a meiotic chromosome axis factor, <u>REV7</u>, and <u>MAD2</u>, a spindle assembly checkpoint protein (Clairmont and D’Andrea, 2021; de Krijger et al., 2021a). REV7 is an abundant cellular protein and is unique among HORMA proteins, both in its large number of binding partners and in its involvement in multiple distinct pathways. Germline biallelic mutations in the REV7 gene can cause the inherited chromosome instability syndrome, Fanconi Anemia (Bluteau et al., 2016). REV7 adopts the two classic closed and open seatbelt conformations of HORMA proteins, and SHLD3 and REV3 are among its seatbelt dependent binding partners (Clairmont et al., 2020).

REV7 is an important determinant of DNA repair pathway choice (Clairmont and D’Andrea, 2021). When closed REV7 (c-REV7) binds to SHLD3, this interaction promotes the assembly of the Shieldin complex (Findlay et al., 2018; Ghezraoui et al., 2018; Gupta et al., 2018; Tomida et al., 2018). The Shieldin complex in turns blocks DSB end resection, promotes reblunting of the resected DSBs, and promotes NHEJ (Dev et al., 2018; Gao et al., 2018; Mirman et al., 2018; Noordermeer et al., 2018). When c-REV7 binds to REV3 in the POLζ complex, the interaction promotes error-prone Translesion Synthesis (TLS) Repair.

The AAA+ ATPase, TRIP13, along with its substrate adaptor p31^comet^, can open REV7 and release SHLD3 or REV3 (Clairmont et al., 2020; Sarangi et al., 2020). Similarly, TRIP13 and p31^comet^ are known to open other HORMA proteins, such as MAD2 (Brulotte et al., 2017; Miniowitz-Shemtov et al., 2015; Ye et al., 2015). The mechanism by which REV7 is converted from the inactive open conformation back to the active closed form is less well understood, and it may involve either the binding of another, unknown SBM-containing protein or a new post-translational modification.

REV7 has at least one additional demonstrated seatbelt-binding partner, CHAMP1 (also known as C13orf80, CAMP, or ZNF828). CHAMP1 is a little-known but highly conserved zinc finger protein first identified as a REV7 interactor (Itoh et al., 2011). CHAMP1 localizes to chromosomes, recruits REV7 to spindles, and plays a role in kinetochore-microtubule interactions. Disruption of CHAMP1 leads to characteristic defects in chromosome alignment in mitosis. Germline heterozygous mutations in CHAMP1 are associated with a rare syndromic form of intellectual disability in humans (Isidor et al., 2016). Crystallographic analysis of the REV7/CHAMP1 complex (Hara et al., 2017) revealed a strong similarity to the REV7/REV3 and REV7/SHLD3 interaction surface (Hara et al., 2010). Despite the clear role of REV7 in DNA repair pathway choice, little is known about the role of its interactor CHAMP1 in DNA repair.

Here, we demonstrate that the interaction of REV7 and CHAMP1 is required for DSB end resection and error-free HR repair. CHAMP1 binds directly to the seatbelt domain of REV7 and thereby competes with the binding of SHLD3 and REV3. DNA damage and the ATM kinase promote the closing of the REV7 seatbelt, resulting in the increased interaction of REV7 with all three binding partners. High cellular levels of CHAMP1 protein favor HR repair over NHEJ and TLS and are often observed in human tumors with acquired HR proficiency. Moreover, CHAMP1 is the active component of a large, multisubunit heterochromatin complex containing HP1α, LEDGF, HDGFRP2, and POGZ previously shown to promote HR activity (Baude et al., 2016; Clairmont et al., 2020; Daugaard et al., 2012; Nozawa et al., 2010; Vermeulen et al., 2010). One function of this complex is to sequester REV7 away from other error-prone repair pathways, under specific cellular conditions and at specific regions of the genome.

## RESULTS

### REV7/CHAMP1 Complex promotes Homologous Recombination Repair

In order to determine the possible involvement of CHAMP1 in DNA repair, we knocked down CHAMP1 expression with siRNA in U2OS cells (**Figure 1**). Interestingly, CHAMP1 knockdown resulted in a reduction in HR activity, based on the decrease in GFP fluorescence generated by the DR-GFP template versus the EJ5-GFP template (Pierce et al., 1999; Stark et al., 2004) (**Figure 1A, B** and **Figure S1A**). Since an early step in HR repair is double strand break (DSB) end resection (Symington, 2014), we used the SMART assay (Huertas and Cruz-Garcia, 2018) to quantify resection. Indeed, two siRNAs to CHAMP1 decreased DSB end-resection (**Figure 1C**). Cells with an HR deficiency have a defect in RAD51 foci assembly and exhibit sensitivity to PARP inhibitors (Bryant et al., 2005; Farmer et al., 2005). Accordingly, RPE-1 cells or U2OS cells with a CRISPR-Cas9-mediated knockout of CHAMP1 exhibited reduced RAD51 foci (**Figure 1D**) and were sensitive to the PARP inhibitor, olaparib (**Figure 1E, F** and **Figure S1B-D**). Previous studies have demonstrated that CHAMP1 interacts directly with REV7 (Hara et al., 2017), a known regulator of DNA HR repair (Boersma et al., 2015; Xu et al., 2015). To confirm and extend these findings, we showed that DNA damage with high dose UV radiation activates the binding of CHAMP1 and REV7 and stimulates the colocalization of REV7 and CHAMP1 in nuclear foci (**Figure S1E-G**). Moreover, CHAMP1 promotes the chromatin localization of REV7 (**Figure S1H, I).** Taken together, we reasoned that the DNA damage inducible interaction of CHAMP1 and REV7 might be required for HR repair.

**Figure 1.**
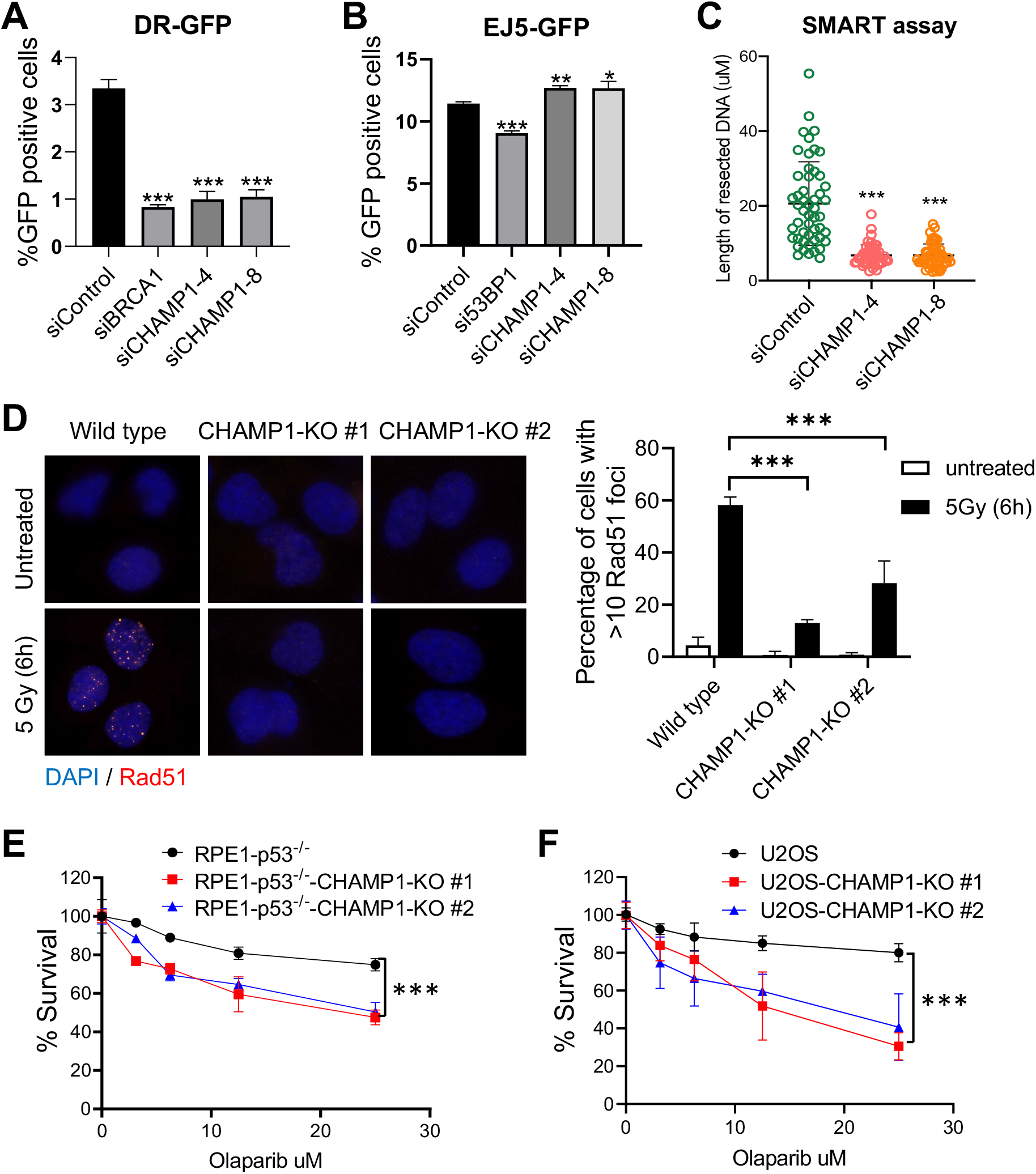
CHAMP1 promotes homologous recombination. **A**, Graph showing the percentage of GFP-positive cells after DR-GFP analysis. U2OS cells were infected with I-SceI adenovirus and knocked down for BRCA1 or CHAMP1 using siRNA. N=3 biologically independent experiments. Error bars indicate standard errors, and p values were calculated using two-tailed Student t-test, ***P<0.0001. **B**, Graph showing the percentage of GFP-positive cells after EJ5-GFP analysis. U2OS cells were infected with I-SceI adenovirus and knocked down for 53BP1 or CHAMP1 using siRNA. N=3 biologically independent experiments. Error bars indicate standard errors, and p values were calculated using two-tailed Student t-test, ***P<0.0001, **P<0.001,*P<0.05. **C**, Quantification of resected ssDNA measured by SMART assay in U2OS cells treated by siControl or siRNAs targeting CHAMP1 for 48hrs. Approximately 50 fibers were counted per experiment. Error bars indicate standard errors, and p values were calculated using Student t-test, ***P<0.0001. **D**, (left) Representative images of RAD51 foci formation in wild-type and two CHAMP1 knockout U2OS cell lines 6 hours after 5Gy IR treatment. (right) Quantification of >10 RAD51 foci. n=3 biologically independent experiments. ***P < 0.001. Statistical analysis was performed using two-tailed student’s t-tests. **E**, 5-day cytotoxicity analysis of wild type and two CHAMP1 knockout RPE1(p53-/-) cell lines treated with various doses of Olaparib; n=3 independent experiments. Wild type versus CHAMP1-KO#1, ***P<0.0001; Wild type versus CHAMP1-KO#2, ***P<0.0001; statistical analysis was performed using two-way ANOVA. **F**, 3-day cytotoxicity analysis of wild type and two CHAMP1 knockout U2OS cell lines treated with various doses of Olaparib. Cell viability were detected by CellTiter-Glo (Promega)n=3 independent experiments. Wild type versus CHAMP1-KO#1, ***P<0.0001; Wild type versus CHAMP1-KO#2, ***P<0.0001; statistical analysis was performed using two-way ANOVA.

### DNA damage activates REV7 seatbelt closure and partner protein binding

We next determined the mechanism by which DNA damage activates the closing of the REV7 seatbelt (**Figure 2**). REV7 has a single, highly-conserved TQ site (T103) which is a possible site of DNA damage-inducible, ATM-dependent phosphorylation (**Figure S2A**) (Matsuoka et al., 2007). Interestingly, this TQ site aligns with a negatively-charged amino acid (E105) in the primary sequence of another HORMA protein, MAD2. Moreover, in the closed conformation of MAD2, an electrostatic interaction between E105 and the positively-charged K192, likely contributes to the closing of the MAD2 seatbelt (**Figure S2B**). Similarly, REV7 has a K198 residue at the corresponding site. We therefore reasoned that a DNA-damage inducible, ATM-dependent, phosphorylation of T103 of REV7 could account, at least in part, for the DNA-damage inducible closing of REV7 and the binding of proteins with a SBM, such as SHLD3, REV3, and CHAMP1. To test this hypothesis, we initially determined whether DNA damage activates the phosphorylation of REV7 at T103, using an anti-p(S/T)Q antibody (**Figure S2C**). Indeed, DNA damage activated the phosphorylation of REV7, and a point mutation of REV7 (T103A) reduced this UV-activated phosphorylation of REV7 *in vitro* **(Figure 2A**) and its chromatin recruitment (**Figure 2B**). Consistent with this, an ATM inhibitor reduced the DNA damage-dependent phosphorylation of REV7, reduced the chromatin recruitment of REV7, and decreased the assembly of REV7 foci (**Figure 2C, D** and **Figure S2D, E**). Similarly, UV damage failed to activate the assembly of nuclear foci of the REV7-T103A mutant protein, confirming that REV7 closing correlates with nuclear foci formation (**Figure S2F**). Unlike wildtype REV7, the REV7-T103A mutant protein failed to reduce RAD51 foci (**Figure S2G, H**) and failed to restore PARP inhibitor sensitivity in REV7(-/-) cells (**Figure S2I, J**). Moreover, knockdown of TRIP13 or p31 resulted in increased binding of CHAMP1 to REV7, while overexpression of TRIP13 reduced this interaction, similarly to our previous findings with the other seatbelt interactors SHLD3 and REV3 (**Figure 2E, F**). The REV7-T103A mutant exhibited reduced binding to either SHLD3, CHAMP1, or REV3 (**Figure 2G-I**). Taken together, DNA damage activates the ATM-dependent phosphorylation of T103 on REV7, thereby promoting the closing of the REV7 seatbelt and the binding of SBM proteins, such as SHLD3 and CHAMP1. The TRIP13/p31 complex opens REV7 and releases these binding partners.

**Figure 2.**
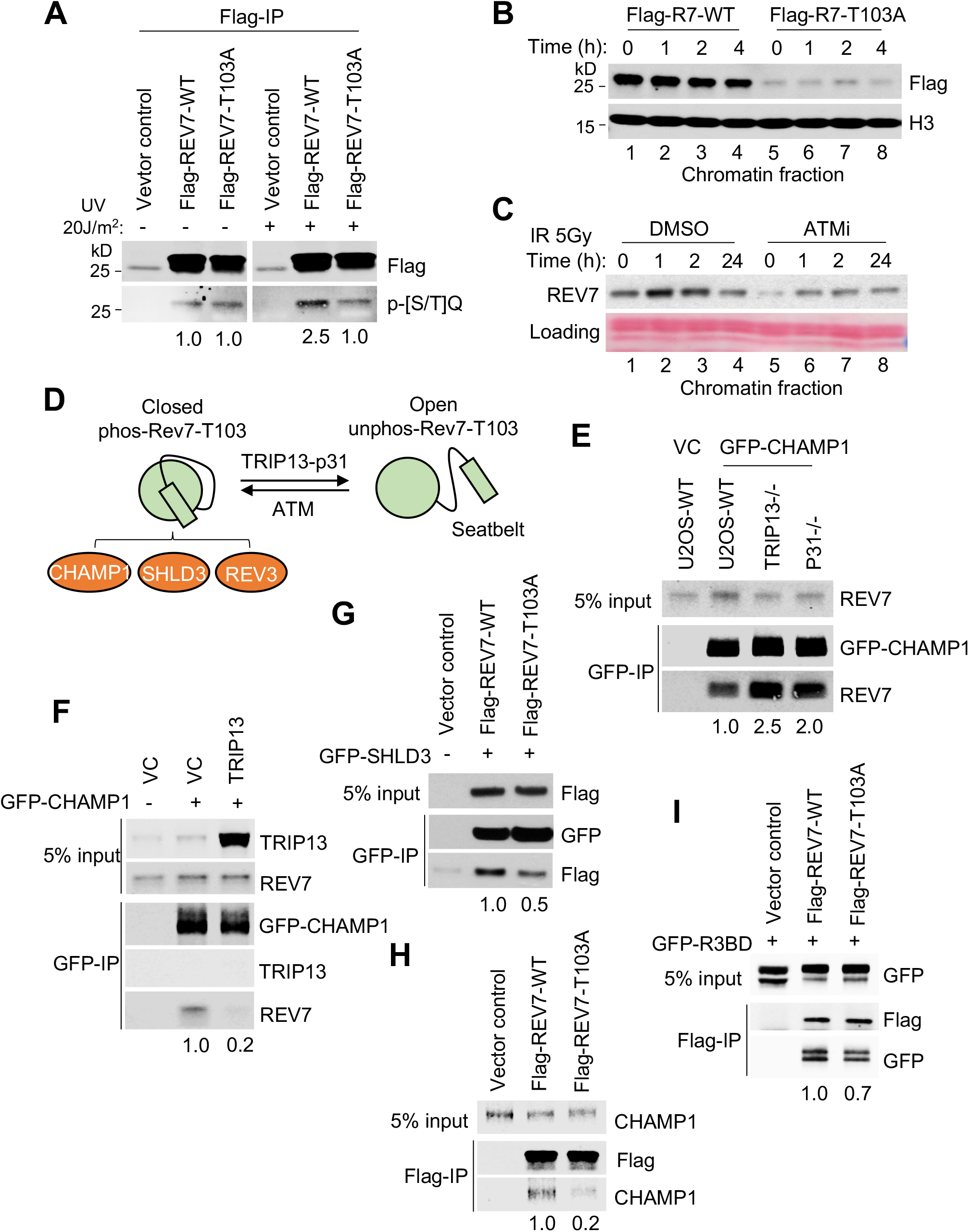
DNA Damage Activates REV7 seatbelt closure and partner protein binding. **A**, 293T cells were transfected with FLAG-REV7-WT or FLAG-REV7-T103A, and following treatment with/without UV (20J/m^2^) for 1 hour. The FLAG-immunoprecipitations were detected by western blot using anti-Flag and anti-p-[S/T]Q antibodies. **B**, 293T cells were transfected with FLAG-REV7-WT or FLAG-REV7-T103A, and following treatment with/without UV (20J/m^2^) as indicated. Western blot showing chromatin fraction of FLAG-REV7-WT and FLAG-REV7-T103A. Histone H3 is used as control for chromatin isolation. **C**, Western blot showing chromatin fraction of REV7 in U2OS treated with DMSO or ATM inhibitor, following IR treatment as indicated. **D**, Schematic of our proposed model showing that the conformational state is regulated by TRIP1-p31 complex and ATM. ATM phosphorylates REV7 at T103 site and promotes the closed form of REV7. The closed REV7 interacts with CHAMP1, SHLD3 and REV3. **E**, Western blot showing GFP-immunoprecipitation of GFP-CHAMP1 in wild-type (WT), *TRIP13^-/-^* and *p31^-/-^* U2OS cells, and the co-immunoprecipitation of endogenous REV7. **F**, Western blot showing GFP-immunoprecipitation of GFP-CHAMP1 in U2OS-vector control (VC) and TRIP13 overexpressed U2OS cells, and the co-immunoprecipitation of endogenous TRIP13 and REV7. **G**, 293T cells were co-transfected with GFP-SHLD3 and Flag-REV7 or Flag-REV7-T103A. Western blot showing GFP-immunoprecipitation of GFP-SHLD3, and the co-immunoprecipitation of Flag-REV7 and Flag-REV7-T103A. **H**, Western blot showing Flag-immunoprecipitation of Flag-REV7 wild type and Flag-REV7-T103A mutant, and the co-immunoprecipitation of endogenous CHAMP1. **I**, 293T cells were co-transfected with GFP-tagged fragment of REV3 containing the REV7-binding domain (R3BD) and Flag-REV7 or Flag-REV7-T103A. Western blot showing Flag-immunoprecipitation of Flag-REV7 and Flag-REV7-T103A, and the co-immunoprecipitation of GFP-R3BD. All of the immunoblots are representative of at least two independent experiments.

### CHAMP1 increases HR activity by competing with SHLD3 for binding to REV7

REV7 is an abundant cellular protein, and it has several known binding partners (Noordermeer et al., 2018). Some of these binding partners bind to the C-terminal seatbelt domain of REV7, including SHLD3, REV3, and CHAMP1 (Clairmont and D’Andrea, 2021; de Krijger et al., 2021a). We reasoned that these partners might compete for seatbelt binding under different cellular conditions or cell cycle stages. The REV7 seatbelt binding protein, SHLD3, promotes the assembly of the Shieldin Complex (Dev et al., 2018; Ghezraoui et al., 2018; Gupta et al., 2018; Noordermeer et al., 2018), thereby blocking the resection of DSBs, recruiting the CST/Polα complex (Barazas et al., 2018; Mirman et al., 2018), and promoting blunt end ligation via the NHEJ pathway. The TRIP13 ATPase, along with its binding partner p31^comet^, opens the seatbelt of REV7 and releases SHLD3 (Clairmont et al., 2020; Sarangi et al., 2020).

As CHAMP1 is much more abundant in cells than SHLD3 (**Figure S3A**), we determined whether CHAMP1 regulates REV7 binding to SHLD3. In HEK293T cells, GFP-SHLD3 binds to REV7, and siRNA knockdown of CHAMP1 resulted in increased co-immunoprecipitation of these proteins, demonstrating that CHAMP1 functions as a negative regulator of the Shieldin complex (**Figure 3A, B**). The REV7 seatbelt also binds to REV3, and the REV7/REV3 (POLζ) complex promotes error-prone Translesion DNA Synthesis (TLS) and enhanced point mutagenesis. We reasoned that CHAMP1 might also sequester REV7 from the REV7/REV3 complex and reduce error-prone TLS activity. To test this hypothesis, we generated and expressed a GFP fusion protein containing the seatbelt binding domain of REV3. As predicted, knocking out CHAMP1 resulted in an increased binding of the TLS polymerase subunit REV3 to REV7 in U2OS cells (**Figure 3C, D**). Consistent with this result, CRISPR-Cas9-mediated knockout of CHAMP1 in U2OS cells or RPE-1 cells resulted in increased REV7/REV3 activity, as measured by MMC resistance and reduced MMC-induced chromosome radials (**Figure 3E, F)** and **Figure S3B-D**). The TRIP13/p31 complex promotes the ATP-dependent opening of the REV7/SHLD3 complex and releases SHLD3 (Clairmont et al., 2020). Taken together, these results support a mechanism in which CHAMP1 promotes DSB end resection by sequestering REV7 from SHLD3 and preventing the assembly of the Shieldin complex.

**Figure 3.**
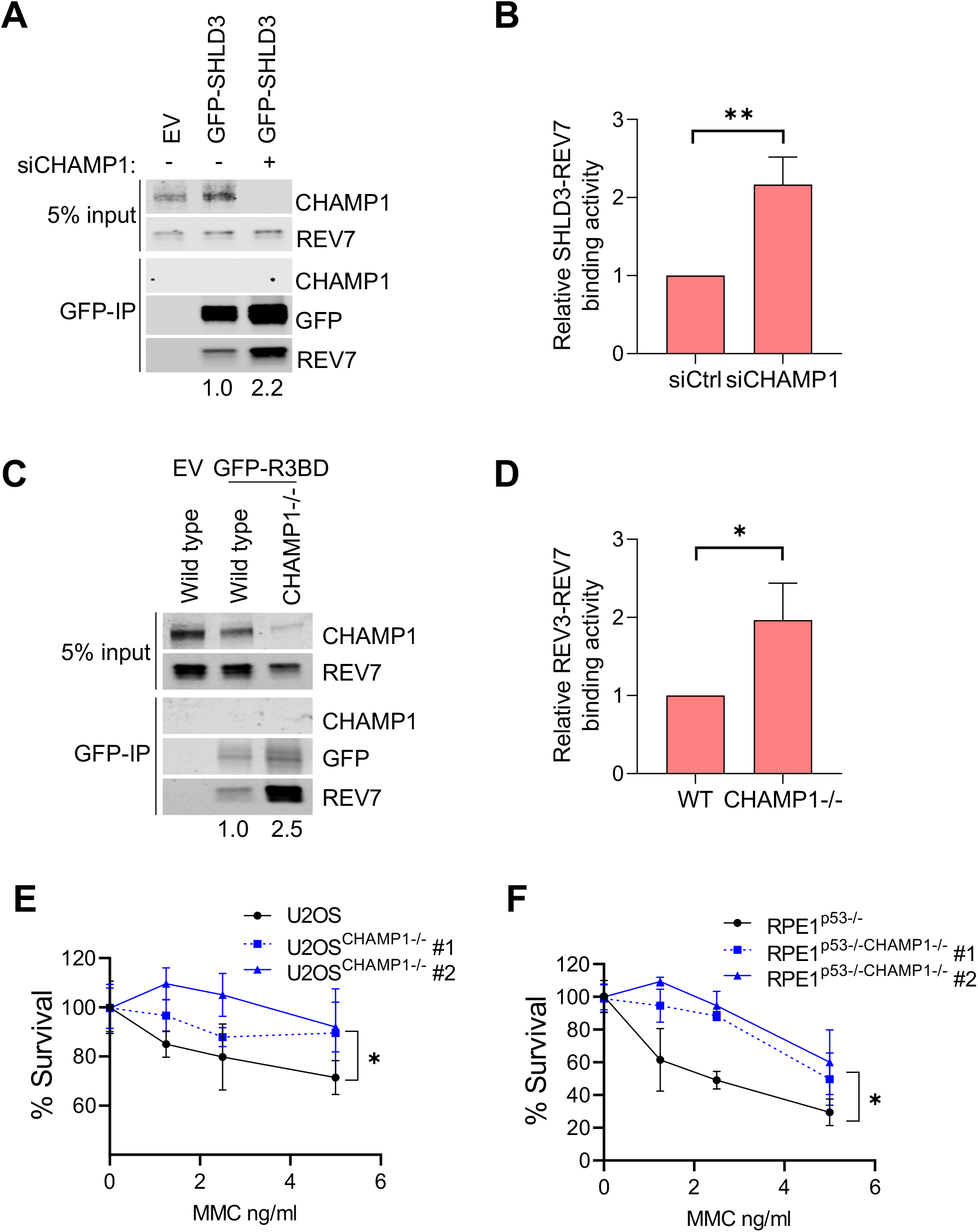
CHAMP1 competes with SHLD3 and REV3 for binding to REV7. **A**, Western blot showing GFP-immunoprecipitation of GFP-SHLD3 in 293T cells, treated with or without siCHAMP1, and the co-immunoprecipitation of endogenous CHAMP1 and REV7. **B**, Quantification of the relative SHLD3-REV7 binding activity from three independent immunoprecipitation western blot shown in A. **C**. Western blot showing GFP-immunoprecipitation of GFP-tagged fragment of REV3 containing the REV7 binding domain (R3BD) in U2OS wild type and U2OS^CHAMP1-/-^ cells, and the co-immunoprecipitation of endogenous CHAMP1 and REV7. **D**, Quantification of the relative REV3-REV7 binding activity from three independent immunoprecipitation western blot shown in C. **E**, A 14 days clonogenic assay of U2OS wild type and two CHAMP1-KO U2OS cell lines, treated with various doses of MMC; n=3 independent experiments. *P<0.05. Statistical analysis was performed using two-way ANOVA. **F**, A 14 days clonogenic assay of RPE1^p53-/-^ and two RPE1^p53-/-CHAMP1-/-^ cell lines, treated with various doses of MMC; n=3 independent experiments. *P<0.05. Statistical analysis was performed using two-way ANOVA.

### The REV7 binding activity of CHAMP1 is required for HR repair but not for proper chromosome alignment

In its primary sequence, CHAMP1 has non-overlapping N-ZNF (C2H2-Zn finger domains), SPE (PxxSPExxK motifs), WK (SPxxWKxxP motifs), FPE (FPExxK motifs), and C-ZNF regions (Itoh et al., 2011) (**Figure 4A**). While the CHAMP1-WK region is required for REV7 binding and recruitment of REV7 to spindles, the CHAMP1-FPE region appears to play an independent role in chromosome alignment (Itoh et al., 2011). To confirm and extend these results, we generated two mutant forms of CHAMP1. According to the molecular structure of the REV7/CHAMP1 complex (Hara et al., 2017), the WKPAKPAPS – motif of CHAMP1, corresponding to the known consensus of a REV7 Seatbelt Binding Motif (SBM), interacts directly with the seatbelt domain of REV7 (**Figure 4B**), albeit with distinct amino acid residue interactions compared to the REV7/REV3 or the REV7/SHLD3 interactions. We therefore generated a mutant form of CHAMP1 which is predicted to disrupt this REV7 binding interaction (ie, the W334A/K335A double mutation, referred to as the CHAMP1-2A mutation). We also generated an in-frame deletion in CHAMP1 (del-FPE CHAMP1), previously shown to be defective in the rescue of chromosome abnormalities in CHAMP1-/- cells (Itoh et al., 2011).

**Figure 4.**
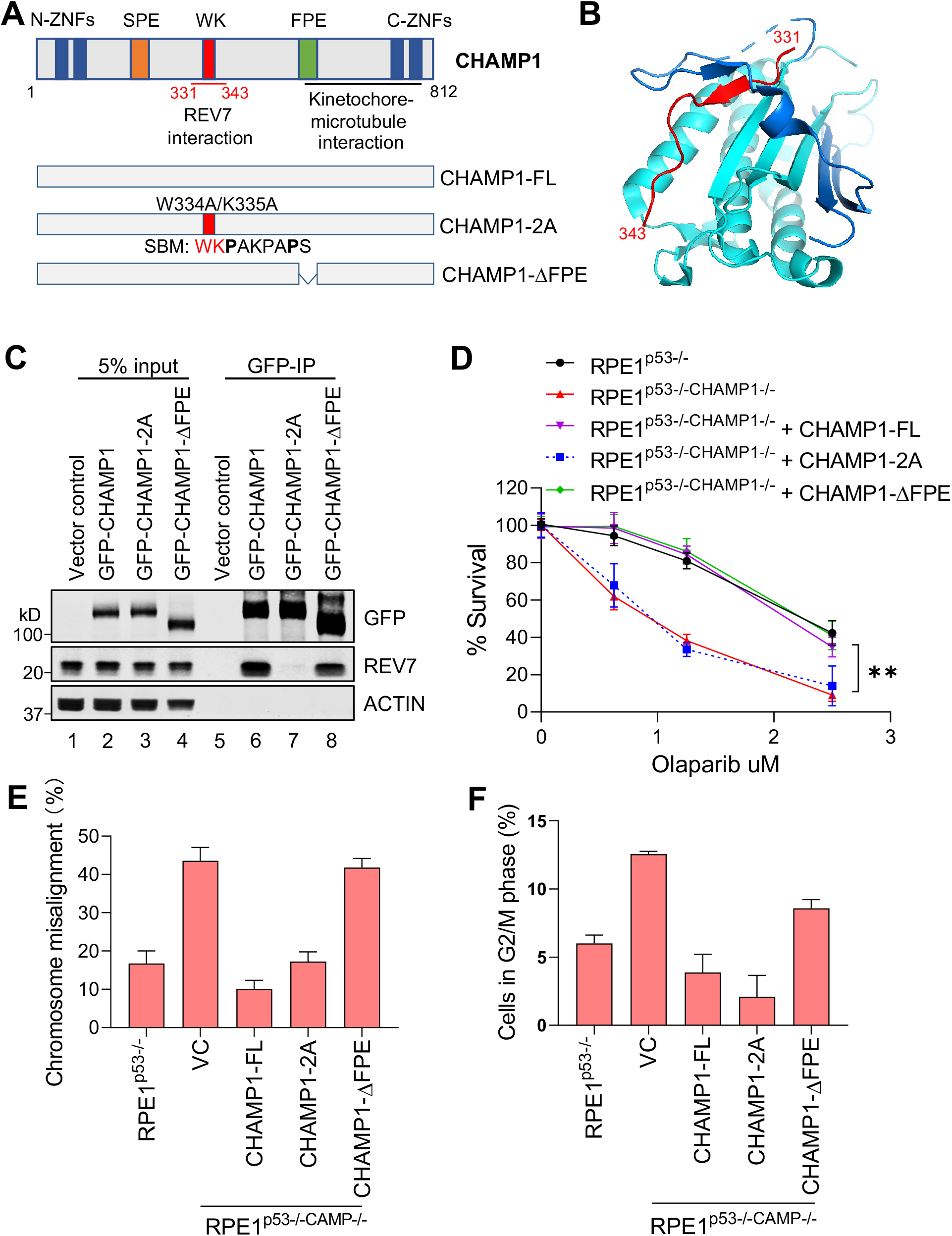
The REV7 binding region of CHAMP1 is required for the HR function but not for correction of chromosome misalignment. **A**, (Top) Schematic of CHAMP1 protein showing its various domains and REV7 binding region. (Bottom) Schematic of CHAMP1-Full Length (FL) and two mutants (2A and ΔFPE). SBM, REV7 seatbelt binding motif. **B**, Structure of the REV7-CHAMP1 complex. REV7 is shown in cyan and blue (seatbelt domain), and the CHAMP1 fragment (residues 331-343) is shown in red. **C**, Western blot showing GFP-immunoprecipitation of GFP-Empty Vector, GFP-CHAMP1 wild-type, GFP-CHAMP1-2A mutant or GFP-CHAMP1-ΔFPE, and the co-immunoprecipitation of endogenous REV7 in 293T cells. **D**, The RPE1^p53-/-CHAMP1-/-^ cells were transfect with vectors containing GFP-tagged CHAMP1-Full Length (FL) or mutants cDNA for 48 hours. GFP positive cells were sorted by Flow Cytometry. A 14 days clonogenic assay of indicated cell lines treated with various doses of Olaparib; n=3 independent experiments, **P<0.001. Statistical analysis was performed using two-way ANOVA. **E**, Summary of chromosome misalignment in indicated cell lines from D. **F**, Same cell lines from D were fixed with 70% ethanol and stained with propidium iodide. Quantitative analysis of indicated cells in G2/M were shown.

As predicted, when expressed in RPE1 CHAMP1-/- cells, the CHAMP1-2A mutant failed to bind to REV7, while the del-FPE CHAMP1 mutant was competent for REV7 binding (**Figure 4C**). Indeed, the CHAMP1-2A mutant failed to correct the PARPi sensitivity of in CHAMP1-/- cells, further confirming that REV7 binding and sequestration by CHAMP1 is required for enhancement of HR activity (**Figure 4D**). The failure of CHAMP1-2A to restore PARP inhibitor resistance and to increase MMC sensitivity was confirmed in the U2OS wildtype or CHAMP1-/- cells (**Figure S4A-E**). Interestingly, complementation with the CHAMP1 del-FPE mutant yielded PARPi resistance, indicating that the FPE domain is not required for enhancement of HR activity.

We next evaluated these two mutant proteins for their ability to correct chromosome misalignment in CHAMP1-/- cells. Consistent with a previous report (Itoh et al., 2011), CRISPR-knockout of CHAMP1 in RPE1 cells results in a severe defect in chromosome alignment (**Figure S4F, G**). The CHAMP1 del-FPE mutant protein failed to complement the chromosome misalignment and the G2/M accumulation of the CHAMP1-/- cells, but the WT CHAMP1 protein or the CHAMP1-2A mutant were functional in these assays (**Figure 4E, F**). Taken together, the WK and FPE domains of CHAMP1 have independent, non-overlapping functions. Moreover, REV7 binding to CHAMP1 is required for HR activity; however, the CHAMP1-mediated recruitment of REV7 to the spindle is not required for the correction of chromosome alignment.

### CHAMP1 regulates homologous recombination through REV7

CHAMP1 therefore regulates HR activity through its ability to sequester REV7 from the Shieldin complex. Thus, we expect that eliminating CHAMP1 in cells already lacking REV7 should not affect HR (**Figure 5**). HR activity can be scored by measuring the level of RAD51 foci or pRPA foci, both known to be increased in the setting of HR. As predicted siRNA knockdown of CHAMP1 in wild-type RPE1 cells reduced the HR activity but failed to reduce the HR activity in cells in which REV7 was already knocked out **(Figure 5A-D**). The REV7-/- cells exhibited increased HR activity, as measured by PARP inhibitor resistance (**Figure 5E**), regardless of their CHAMP1 expression level. Consistent with these results, knockdown of CHAMP1 in a SHLD2-deficient cell line, HCC1937, also failed to cause Olaparib sensitivity **(Figure S5A).**

**Figure 5.**
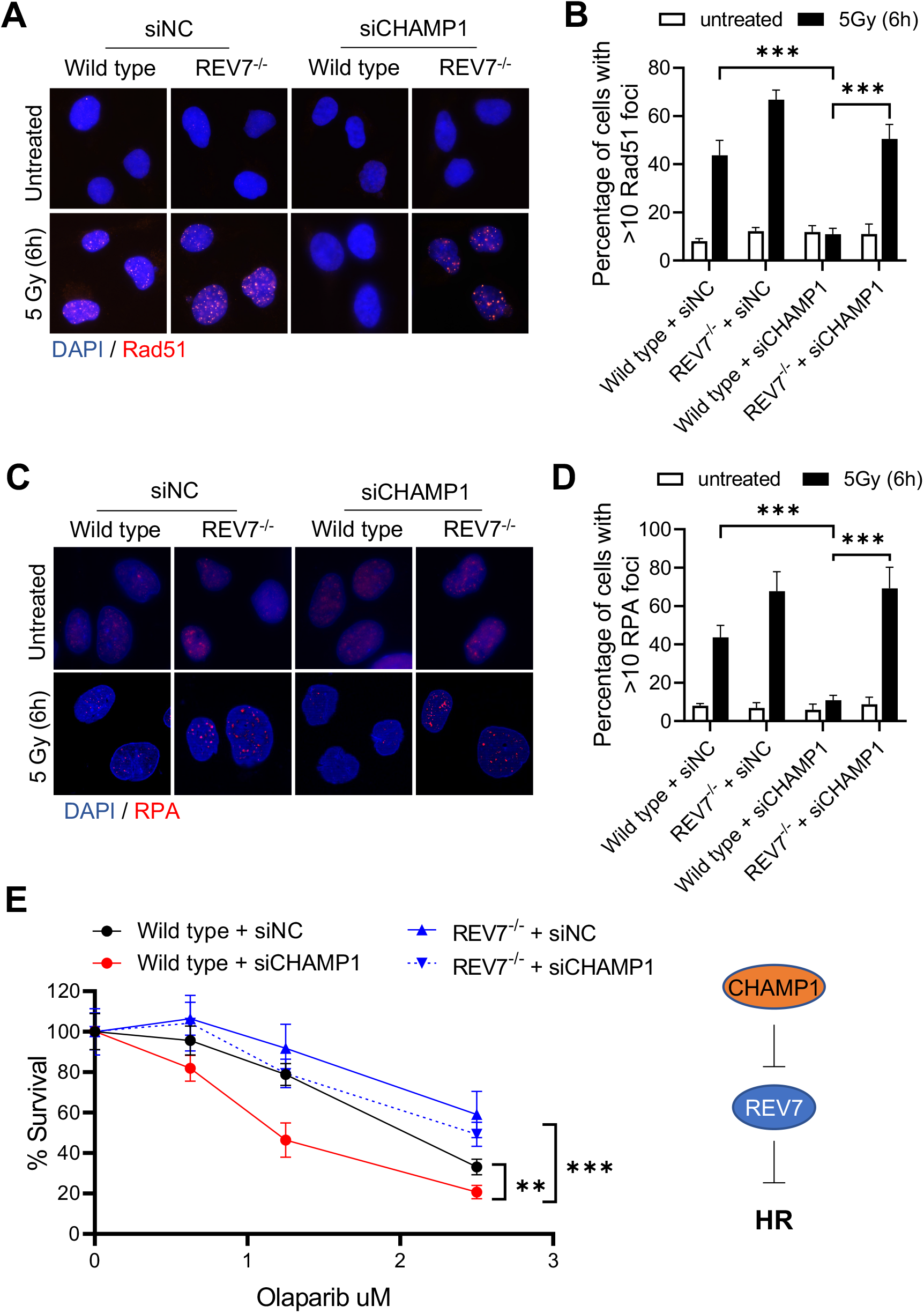
CHAMP1 regulates HR through REV7. **A**, Representative images of RAD51 foci formation in wild-type and *REV7^-/-^* U2OS cells treated with siRNA negative control (siNC) and siCHAMP1, and 6 hours after 5Gy IR treatment. **B**, Quantification of RAD51 in A. More than 10 RAD51 foci were counted. n=3 biologically independent experiments, ***P < 0.0001. Statistical analysis was performed using two-tailed student’s t-tests. **C**, Representative images of p-RPA32(S33) foci formation in wild-type and *REV7^-/-^* U2OS cells treated with siRNA negative control (siNC) and siCHAMP1, and 6 hours after 5 Gy IR treatment. **D**, Quantification of p-RPA32(S33) foci in C. More than 10 RAD51 foci were counted. n=3 biologically independent experiments, ***P < 0.0001. Statistical analysis was performed using two-tailed student’s t-tests. **E**, (left) A 14-days clonogenic survival of wild-type and REV7^-/-^ U2OS cells treated with various doses of Olaparib after siControl or siCHAMP1 treatment. n=3 independent experiments, **P < 0.001, ***P < 0.0001. Statistical analysis was performed using two-way ANOVA. (right) A cartoon shows that CHAMP1 inhibits REV7 to promote HR.

To further validate this model, we next sought clinical evidence that CHAMP1 expression might affect cancer patient survival (**Figure S5B, C**). We reasoned that human tumors with an underlying defect in an HR pathway might upregulate CHAMP1 as a compensatory mechanism to tolerate their low HR and their replication stress. To test this hypothesis, we correlated the level of CHAMP1 expression in ovarian tumors with patient survival. For patients with tumors with low REV7 expression, the level of expression of CHAMP1 did not affect survival (**Figure S5B).** Thus, consistent with the cellular data, the elevated HR activity in cells with low or absent REV7 expression was unaffected by CHAMP1 expression levels. Interestingly, for patients with tumors with high REV7 expression, the level of CHAMP1 expression significantly affected patient survival (**Figure S5C)**. The high CHAMP1 expression correlated with a more aggressive tumor and poor patient prognosis, perhaps resulting from the improved HR activity of these tumors. Taken together, the ability of CHAMP1 to enhance HR is directly dependent on the presence of the REV7 protein.

### CHAMP1 overexpression is common in tumors with underlying HR deficiency and correlates with poor cancer patient prognosis

We next sought additional evidence that CHAMP1 upregulation correlates with PARP inhibitor resistance. We used a panel of BRCA1-deficient cell lines with acquired PARPi-resistance, collected through serial selection in increasing concentrations of PARPi (**Figure 6A**) (Farkkila et al., 2021). These cells exhibited multiple independent mechanisms of PARPi resistance, including downregulation of the Shieldin Complex or upregulation of ATR/CHK1 pathway activity (Farkkila et al., 2021). Interestingly, one of these PARPi-resistant clones (NA5) exhibited high CHAMP1 protein expression compared to the parental PARPi-sensitive cell line (**Figure S6A**). Knockdown of CHAMP1 in these cells restored PARPi sensitivity (**Figure 6B**). In contrast, knockdown of CHAMP1 in another line (NA1), which has a lower level of CHAMP1, did not restore PARPi sensitivity. Taken together, *BRCA1*-deficient cells can acquire PARPi resistance, at least in part, by upregulating CHAMP1 expression.

**Fig 6.**
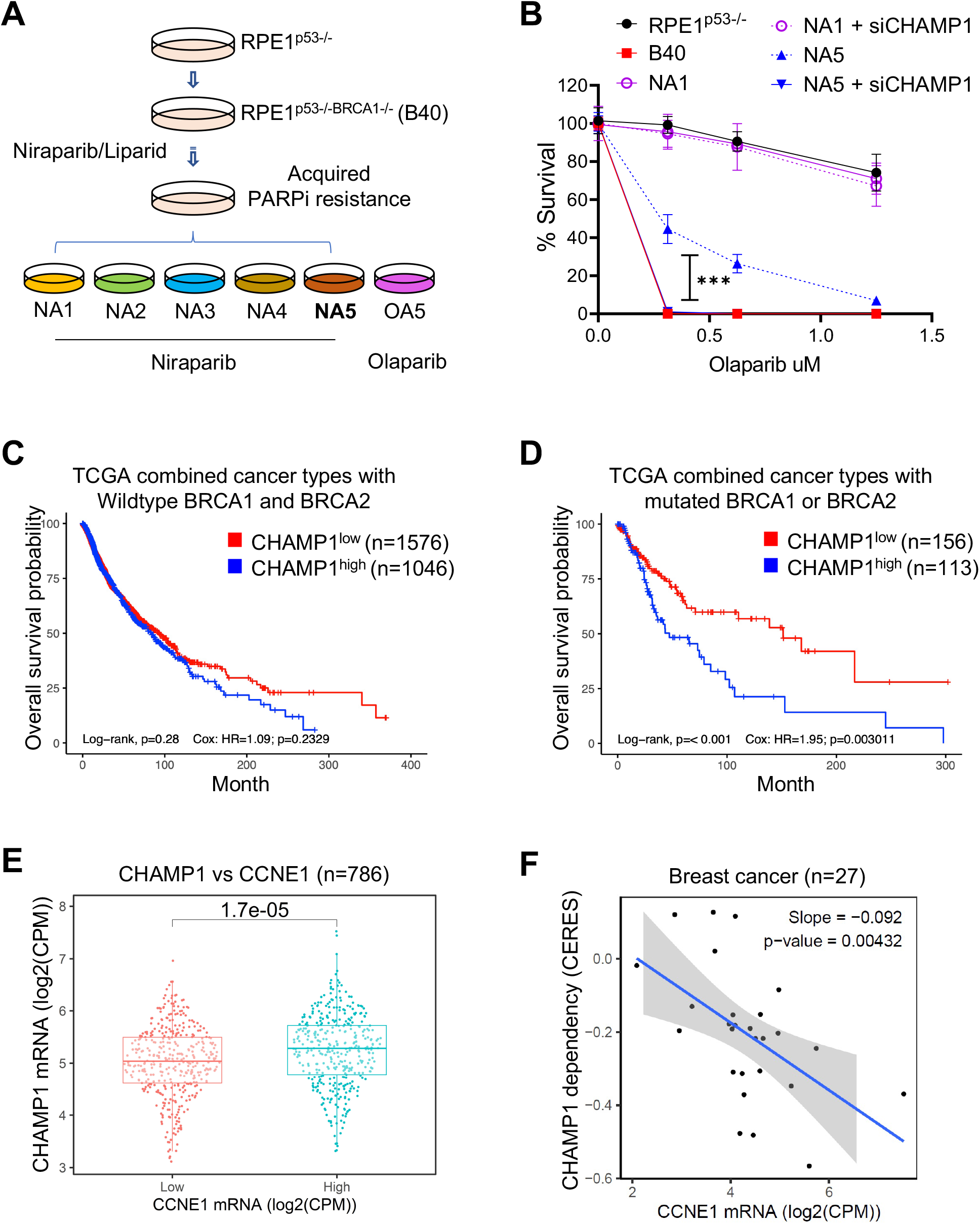
CAMP overexpression is common in tumors with underlying HR deficiency and correlates with poor cancer patient prognosis. **A**, Schematic of PARPi-resistant RHE1^p53-/-BRCA1-/-^ cells generation. RPE1P^53-/-BRCA1-/-^ cells (B40) were treated with increasing concentrations of the PARPis niraparib/Olaparib over 3 months, and then isolated by single-cell clones from the niraparib- and Olaparib-resistant pools. **B**, A 14-days clonogenic survival of RPE1^p53-/-^, RPE1^p53-/-BRCA1-/-^ and niraparib/Olaparib-resistant RPE1^p53-/-BRCA1-/-^ cell clones treated with various doses of Olaparib after siControl or siCHAMP1 treatment. n=3 independent experiments. ***P < 0.0001. Statistical analysis was performed using two-way ANOVA. **C-D**, Kaplan–Meier curves depicting overall survival of patients from TCGA with CHAMP1 expression and wildtype BRCA1 and BRCA2 (**C**), and mutated BRCA1 or BRCA2 (**D**). This analysis combines tumors from these TCGA studies: BLCA (blader), BRCA (breast), LUAD (lung), LUSC (lung squamous), and SKCM (skin). **E**, CHAMP1 expression positively correlates with Cyclin E expression. **F**, Breast cancer cells with high expression of CCNE1 are more dependent on CHAMP1 for survival.

Additional analysis of clinical databases revealed that, for patients with HR deficient tumors containing a BRCA1 or BRCA2 mutation, a high level of CHAMP1 expression correlates with a worse prognosis (**Figure 6C, D**). This result further suggests that high CHAMP1 expression can partially correct the HR deficiency of these tumors, leading to a more aggressive tumor phenotype. Tumors with BRCA2 mutations and presumably HR deficiency exhibited higher baseline levels of CHAMP1 mRNA expression (**Figure S6B**). Consistent with these observations, the level of CHAMP1 mRNA expression in cancer cell lines strongly correlates with the level of CCNE1 mRNA expression (**Figure 6E**). Cells with a high degree of replication stress resulting from CCNE1 amplification may therefore rely on CHAMP1-mediated HR for their survival. Indeed, breast cancer cell lines with high expression of CCNE1 mRNA are more dependent on CHAMP1 for their proliferation and survival (**Figure 6F**).

### POGZ binds to CHAMP1 and cooperates in HR Repair

Recent studies have shown that CHAMP1 is a subunit of a large multisubunit complex of HP1α heterochromatin binding proteins. This complex includes HP1α, POGZ, LEDGF, and HDGFRP2 (Baude et al., 2016; Clairmont et al., 2020; Daugaard et al., 2012; Nozawa et al., 2010; Vermeulen et al., 2010). REV7 coimmunoprecipitates with multiple components of this complex (Noordermeer et al., 2018), further suggesting a functional link with DNA repair regulation. Interestingly, knockdown or knockout of many of the subunits of this complex, such as HP1α (Soria and Almouzni, 2013), LEDGF (Daugaard et al., 2012) or HDGFRP2 (Baude et al., 2016), similar to the knockdown of CHAMP1, reduces DSB end resection and HR and increases PARP inhibitor sensitivity (Olivieri et al., 2020). To confirm and extend these studies, we next evaluated the POGZ subunit of this complex. Knockout of POGZ in RPE cells, like knockout of CHAMP1, resulted in decreased HR repair, decreased DSB end resection, decreased RAD51 foci, and increased PARP inhibitor sensitivity (**Figure 7A-C and Figure S7A,B**).

**Figure 7.**
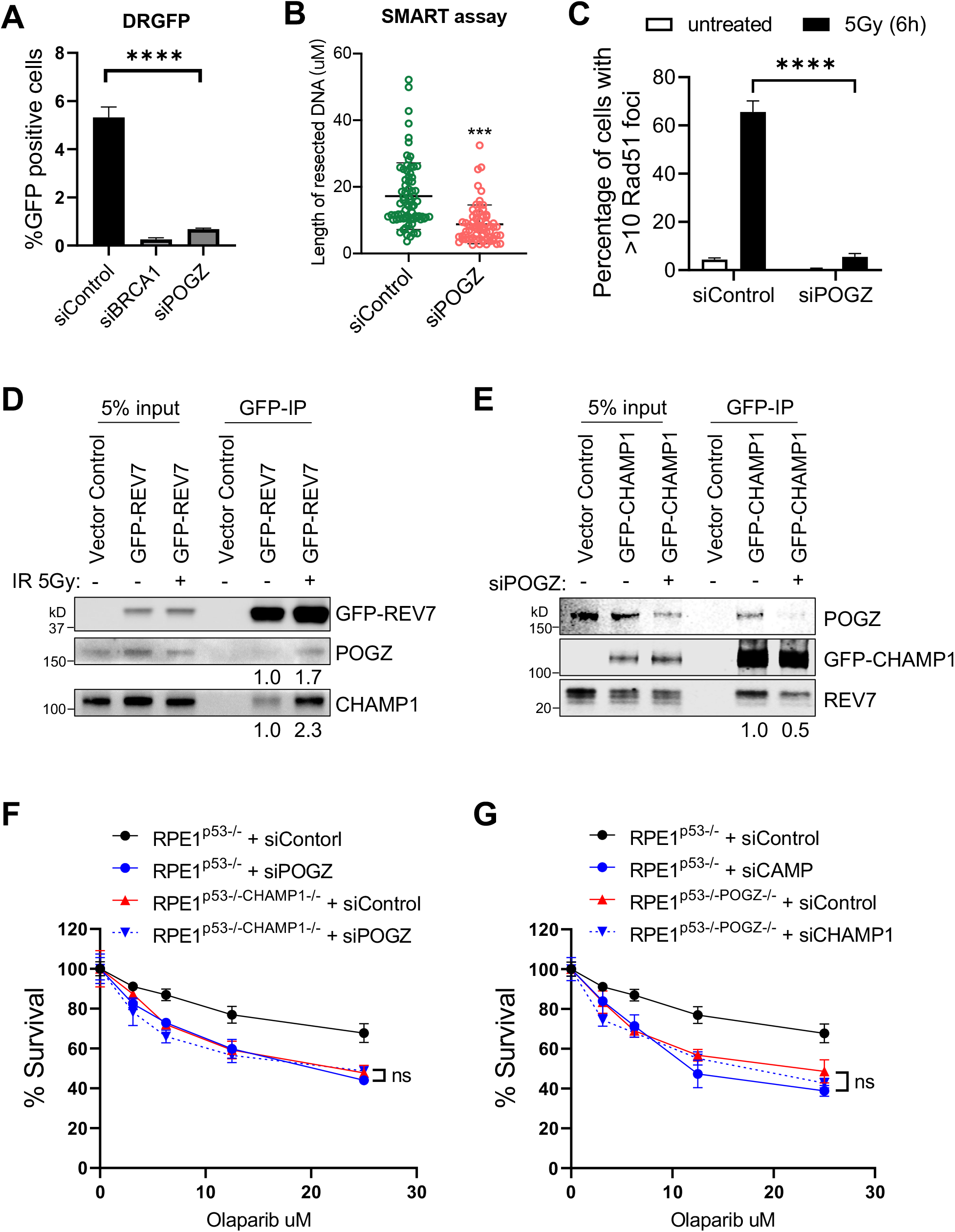
POGZ is epistatic with CHAMP1 in the Regulation of Homologous Recombination. **A**, Graph showing the percentage of GFP-positive cells after DR-GFP analysis. U2OS cells were infected with I-SceI adenovirus and knocked down for BRCA1 or POGZ using siRNA. N=3 biologically independent experiments. Error bars indicate standard errors, and p values were calculated using two-tailed Student t-test, ***P<0.0001. **B**, Quantification of resected ssDNA measured by SMART assay in U2OS cells treated by siControl or siRNAs targeting CHAMP1 for 48hrs. Approximately 50 fibers were counted per experiment. Error bars indicate standard errors, and p values were calculated using Student t-test, ***P<0.0001. **C**, Quantification of >10 RAD51 foci in wild-type and two CHAMP1 knockout U2OS cell lines 6 hours after 5Gy IR treatment. n=3 biologically independent experiments. ***P < 0.001. Statistical analysis was performed using two-tailed student’s t-tests. **D**, Western blot showing GFP-immunoprecipitation of GFP-REV7 in HEK293T cells treated with or without irradiation (5Gy), and the coimmunoprecipitation of endogenous CHAMP1 and POGZ. **E**, Western blot showing GFP-immunoprecipitation of GFP-CHAMP1 in HEK293T cells treated with or without siPOGZ, and the co-immunoprecipitation of endogenous CHAMP1 and POGZ. **F**, 3-day cytotoxicity analysis of RPE1^p53-/-^ and RPE1^p53-/-CHAMP1-/-^ cells treated with various doses of Olaparib after 48 hours siControl or siPOGZ treatment. Cell viability were detected by CellTiter-Glo (Promega), n=3 independent experiments. Statistical analysis was performed using two-way ANOVA. **G**, 3-day cytotoxicity analysis of RPE1^p53-/-^ and RPE1^p53-/-POGZ-/-^ cells treated with various doses of Olaparib after 48 hours siControl or siCHAMP1 treatment. Cell viability were detected by CellTiter-Glo (Promega), n=3 independent experiments. Statistical analysis was performed using two-way ANOVA. All of the immunoblots are representative of at least two independent experiments.

Moreover, IR-induced DNA damage activated the co-immunopreciptation of POGZ and CHAMP1 (**Figure 7D**). Knockdown of POGZ reduced the interaction of REV7 and CHAMP1 (**Figure 7E**), and POGZ binding to CHAMP1 was independent of REV7 seatbelt binding (**Figure S7C**). Importantly, knockdown of POGZ in CHAMP(-/-) cells or knockdown CHAMP1 in POGZ(-/-) cells resulted in no additional impairment of HR repair or PARP inhibitor sensitivity (**Figure 7F, G**), demonstrating that POGZ and CHAMP1 are epistatic in HR repair. Finally, POGZ expression, like CHAMP1 expression, is increased in many human cancers, consistent with its compensatory role in promoting HR repair (**Figure S7D**). Taken together, these results demonstrate that a CHAMP1-containing multisubunit complex has a functional role in sequestering REV7, preventing its association with SHLD3 in the Shieldin complex, and promoting HR repair locally in heterochromatin.

## DISCUSSION

Our results demonstrate that REV7, through its C-terminal seatbelt, can bind to three different factors, SHLD3, REV3, and CHAMP1, to elicit distinct DNA repair outcomes. The REV7-SHLD3 interaction mediates the assembly and accrual of the Shieldin complex at DSBs to block DSB end resection and channel repair through NHEJ. The REV7-REV3 interaction promotes the Polζ Translesion Synthesis (TLS) complex through its interaction with REV1, and this complex bypasses bulky adducts of DNA during replication, thereby promoting mutagenesis. Finally, the REV7-CHAMP1 complex, by sequestering REV7 from REV7-SHLD3 and REV7-REV3 complexes, can promote error-free HR repair and function as a negative regulator of the error-prone NHEJ and TLS repair pathways, respectively. Of the three binding proteins, CHAMP1 is the most abundant, resulting in a trend toward error-free DNA repair. The mechanisms by which cells switch from one REV7 binding complex to another is largely unknown.

Although CHAMP1 may have additional, independent roles in HR repair, its sequestration of REV7 appears to be its primary mechanism in HR activity. Indeed, cells with a knockout of REV7, versus cells with a double knockout of REV7 and CHAMP1, have equally high levels of DSB end resection, RAD51 foci, and PARP inhibitor resistance. These results argue for an epistatic relationship of REV7 and CHAMP1 in HR repair.

Previous studies indicate that TRIP13/p31^comet^ complex opens REV7 and, like CHAMP1, also reduces the Shieldin complex and promotes DSB end resection and HR repair (Clairmont et al., 2020; de Krijger et al., 2021b). TRIP13 and CHAMP1 appear to be non-epistatic in HR repair, however, since knockdown of TRIP13 in CHAMP1 (-/-) cells results in a further decline in HR repair. Accordingly, a reduction in REV7/SHLD3 levels, by either TRIP13/p31^comet^ or CHAMP expression, appears to result from independent mechanisms for upregulating HR repair.

We have shown that DNA damage following ionizing radiation activates the ATM-dependent closing of REV7 and the interaction of REV7 with either SHLD3, REV3, or CHAMP1. How and when REV7 selectively chooses one binding partner versus another is mostly unknown. It will be important to determine whether specific kinds of DNA damage will preferentially activate a specific complex. For instance, IR may preferentially activate the REV7/SHLD3 complex while replication fork perturbants, like hydroxyurea, may activate the REV7/CHAMP1 complex. Also, the choice of a specific REV7 complex may be strongly influenced by either the local organization of the genome, cell cycle cues, or cell type specificity. The specific interaction of REV7 with SHLD3, REV3, or CHAMP1 may also be determined by the TRIP13/p31^comet^ enzyme. While all three complexes are released by TRIP13/p31^comet^, cells may choose to selectively release one complex or another, depending on specific cellular demands for NHEJ, TLS, or HR, respectively.

In addition to its role in the Shieldin complex and upregulation of NHEJ repair, REV7 is also a critical determinant of Translesion DNA Synthesis (TLS repair). Indeed, one of the best known roles of REV7 is its role as the small non-catalytic subunit of the Polζ complex. Pol ζ is one of several human polymerases, specialized for synthesizing DNA across lesions in the template strand, a process known as translesion synthesis (TLS). TLS is far more mutagenic than normal replication (Goodman and Woodgate, 2013; Sale, 2013). In particular, DNA POLζ, in conjunction with its partner, REV1, are responsible for the majority of spontaneous and damage induced mutations during DNA replication (Gibbs et al., 2000; Jansen et al., 2005).

Although the mutagenic TLS and NHEJ pathways have historically been viewed as independent processes, their coordinate regulation by REV7 calls for a reevaluation of this relationship. Indeed, there are several important relationships between these processes aside from REV7, and some of these relationships are functional or spatiotemporal. First, in both contexts, the active REV7/REV3 or REV7/SHLD3 complexes promote the activity of rapid mutagenic pathways, in contrast with the slower process of HR repair. Second, both TLS and NHEJ can still play an important role during S-phase when the bulk of DNA repair is carried out HR. Lastly, resection and the HR pathway are well-known to be utilized at stalled replication forks as well as DSBs, raising the possibility that REV7/REV3 and REV7/SHLD3 could act on the same substrate.

Finally, additional factors may influence the relative levels of REV7/SHLD3, REV7/REV3, and REV7/CHAMP1 in the cell. First, REV7 may preferentially bind to CHAMP1 in heterochromatic regions of the genome, such as centromeres and telomeres, resulting in higher local levels of HR repair. Second, CHAMP1 may also have a distinct binding affinity for the REV7 seatbelt. Indeed, based on the corresponding crystal structures, the molecular interactions of the REV7 seatbelt with either CHAMP1 or REV3 are distinct (Hara et al., 2010; Hara et al., 2017) and likely to result in distinct binding affinities and off rates. Third, some tumor cells with an underlying genetic deficiency in HR repair, such as a BRCA1 or BRCA2 mutation, have higher level of expression of CHAMP1. Interestingly, this increase in CHAMP1 may provide these cells with a compensatory increase in HR and a higher capacity for tolerating replication stress. Finally, the specific interaction of REV7 with these various binding partners may be highly regulated by post-translational modifications and under distinct cellular conditions. Future studies are needed to further assess the spatial and temporal control of the REV7 interaction with CHAMP1, SHLD3, and REV3.

## ACKNOWLEDGMENTS

We thank all members of the D’Andrea laboratory for their helpful suggestions and comments. This work was supported by grants from the US National Institutes of Health (R37HL052725 and P01HL048546), the US Department of Defense (BM110181), the Breast Cancer Research Foundation, the Fanconi Anemia Research Fund, the Ludwig Center at Harvard, and the Smith Family Foundation (to A.D.D.) and the Claudia Adams Barr Program in Innovative Basic Cancer Research (to F.L. and P.S).

## AUTHOR CONTRIBUTIONS

F.L., C.C., P.S., and A.D.D. conceived the study, analyzed the data, and wrote the manuscript. H.F., L.M, and H.N. performed experiments and analyzed the data.

## Materials and Methods

### Cell culture and transfections

Human U2OS, RPE1-hTERT, HCC1937 and HEK293T cells were cultured in DMEM/F12 + Glutamax (Invitrogen) supplemented with 10% FBS (Sigma) and 1% penicillin-streptomycin (Invitrogen). DNA transfections and siRNA knockdowns were carried out using Lipofectamine LTX (Invitrogen) and RNAiMax (Invitrogen) respectively according to the manufacturer’s protocols. The individual siRNAs used are: AllStar negative siControl (1027281); siCHAMP1 #4 (SI00973084); siCHAMP1 #8 (SI04282159); siBRCA1 (SI00930510); si53BP1 (SI01456539) were purchased from Qiagen. ON-target Human siPOGZ (L-006953-01-0005) were purchased from Horizon Discovery.

### Antibodies and chemicals

Antibodies used in this study were: Abnova H00283489-B01P (C13orf8/CHAMP1, IB, IF), Abcam ab180579 (Mad2L2/REV7, IB, IF), Bethyl Laboratories A302-509A (POGZ, IB, IF), Abcam ab128171 (TRIP13, IB), Cell Signaling 3873 (alpha-Tubulin, IF), Cell Signaling 2187 (Phospho-CENP-A, IF), Cell Signaling 3638 (H3, IB), Cell Signaling 2956 (GFP, IB), Cell Signaling 3700 (Actin, IB), Abcam ab70369 (phospho-Kap1-S824, IB), Cell Signaling 6966 (Phospho-[S/T]Q, IB), Fisher Scientific NB100544 (RPA2-P-Ser33, IF), Santa Cruz sc-8349 (RAD51, IF), Millipore-Sigma F3165 (GAPDH, IB) and Millipore-Sigma F3165 (Flag, IB, IF). Mitomycin C (MMC) was purchased from Sigma and Olaparib was purchased from Selleckchem.

### Generation of knockout cell lines with CRISPR-Cas9

CHAMP1 and POGZ guide RNA sequences were cloned into the pSpCas9 BB-2A-GFP (PX458) vector (GenScript). U2OS and RPE1^p53-/-^ cells were transfected with Cas9-gRNA plasmids. After 48 hours GFP positive cells were selected using a BD FACSAria II cell sorter. Single cells from GFP positive pool were cultured for three to four weeks and colonies were screened for knockouts by western blotting using the anti-p31^comet^ antibody (Millipore-Sigma). The guide RNA sequences targeting CHAMP1 in this study were: #1 TCGTAAACCATCAGCACGTT and #2 CCAGAGATCCGTAGTCCAGC. The guide RNA sequences targeting POGZ in this study were: #1 CAGTTTGTTAAGCCGACAGT and #2 TCTGCTGATCGAGTTCTACG.

### GFP-based DNA Repair Assays

For DR- and EJ5-GFP reporter assays, U2OS cells carrying the respective GFP expression cassette were transfected with the indicated siRNAs. 24 hours after transfection, cells were infected with or without I-SceI lentivirus. After 48 hours, cells were harvested and detected by flow cytometry. The data was analyzed using the FlowJo software.

### Cellular fractionation and immunoblot analysis

Cells were lysed with NP40 buffer (1% NP40, 300 mM NaCl, 0.1 mM EDTA, 50 mM Tris (pH 7.5)) supplemented with phosphatase and protease inhibitor cocktail (Roche). Cell lysates were resolved by NuPAGE 4-12% Bis-Tris gels (Invitrogen), and transferred onto nitrocellulose membranes. Membranes were blocked with 5% BSA in TBST and were sequentially incubated with primary and secondary antibodies and detected using chemiluminescence or fluorescence (LI-COR Biosciences). For chromatin extraction, chromatin-bound extracts were got using subcellular protein fractionation kit (Thermo). The band intensities were measured by ImageJ.

### Immunoprecipitation

After transfection for 48h, 293T or U2OS cells were then harvested and lysed in NETN lysis buffer with proteinase & phosphatase inhibitor cocktail (Thermo, 1:100) for 30 minutes on ice. They were then incubated with antibody-bead conjugate (Anti-FLAG^®^ M2 Magnetic Beads, Millipore & Sigma or GFP-Trap_A, Chromotek) overnight at 4 °C. Beads were washed four times with NETN buffer and immunoprecipitates were eluted by boiling. Western blots were performed to detect the immunoprecipitates. The band intensities were measured by ImageJ.

### Drug sensitivity assays

Cells were transfected with plasmid or siRNA 24h before being plated for colony formation or CellTiter-Glo assays. To assay clonogenic survival, cells were seeded at 500-1000 cells/well in 6-well plates in triplicates. Drugs at the shown doses were added after 12 hours and cells were permitted to grow for 14 days. Colony formation was scored by fixing and staining with 0.5% (w/v) crystal violet in 20% methanol. For short term CellTiter-Glo survival assays, cells were plated in 96-well plates at 800-1000 cells/well, and treated with drugs at the indicated concentrations after 12 hours. Three days later, cellular viability was measured using CellTiter-Glo (Promega). Survival at each drug concentration was calculated as a percentage normalized to the corresponding untreated control, for both assays.

### Immunofluorescence assays

Cells were plated on glass coverslips in 12-well plates. They were then either left untreated or treated at 20J/m^2^ UV or 5Gy IR. After 1 or 6 hours, they were harvested by pre-extraction with 0.5% Triton X-100 for 5 min, followed 4% paraformaldehyde fixation for 10 min at 4 °C. After three PBS washes, blocking was performed with 3% BSA in PBS for 1 hour at room temperature, followed by sequential primary and secondary antibody incubations overnight at 4 °C and 1 hour at room temperature respectively. The coverslips were mounted with DAPI (Vector Laboratories) and captured using a Zeiss AX10 fluorescence microscope and Zen software, and foci were scored. At least 100 cells were counted for each sample.

### SMART assay

The SMART DNA fiber assay procedure was performed largely as described previously (Clairmont et al., 2020). In brief, cells were treated with BrdU (sigma) for 24h, and then exposed to X-ray irradiation to induce DSB formation. Cells were collected 6h after irradiation, and embedded in low melting point agarose plugs before lysis with proteinase K overnight at 50°C. The plugs were then washed with TE buffer and digested with agarase (NEB). The sample solution was spread onto silanized coverslips using the FiberComb machine (Genomic Vision). Combed coverslips were blocking with 3% BSA for 30min, and then incubated with anti-BrdU antibody (rat, abcam) overnight at 4°C. After incubation with secondary Alexa-555-labelled goat anti-rat antibodies, the coverslips were washed and mounted with Vectashield mounting medium (Vector laboratories). Images were captured by Zeiss AX10 flurorescence microscope. At least 100 fibers were counted per condition. The fiber lengths were measured using imageJ and graphed.

### Chromosomal aberration analysis

RPE1^p53-/-^ and RPE1^p53-/-CHAMP1-/-^ cells were incubated with or without 20 ng/ml MMC for 48 hours. Cells were treated with 100 ng/ml of colcemid for 2 hours, followed by a hypotonic solution (0.075 M KCl) for 20 min and fixed with 3:1 methanol/acetic acid. After staining with Wright’s stain, 50 metaphase spreads were counted for aberrations. The relative number of chromosomal dicentrics and radials was calculated relative to control cells as indicated.

### TCGA data acquisition and analysis

The survival analyses of the Cancer Genome Atlas (TCGA) patients were performed using the clinical and RNASeq expression and genomic alteration data of TCGA Pan-Cancer study for 32 cancer types downloaded from the cBioPortal for Cancer Genomics (https://www.cbioportal.org; retrieved March 2020). For the survival analysis with mRNA expression of CHAMP1 (CHAMP1) and REV7 (MAD2L2), for each cancer type, samples were grouped into the low- and high-mRNA expressing groups for CHAMP1 and REV7 based on the expression z-scores of either zero, or less than −0.5 and greater than 0.5. These expression z-scores were computed relative to the diploid samples. Survival analysis was then performed in R for each cancer type to determine whether there was a difference in the overall survival between the two groups, separately for CHAMP1 and REV7, and for REV7 in each of the two CHAMP1 groups. Kaplan-Meier curves were created, and the log-rank test was used to test for a difference in overall survival using the survival package in R. The p values were calculated from the chi-square distribution. The survminer R package was used to estimate median survivals, and to plot the Kaplan-Meier curves. Additionally, Cox proportional hazards regression was performed to estimate the hazard ratio between the low- and the high-mRNA groups for each cancer type.

The survival analyses of TCGA patients with CHAMP1 mRNA expression and mutation status of BRCA1 and BRCA2 were performed as follows. A tumor was considered mutated for a gene if it had variants with classifications that were damaging or other non-conserving. The analyses were first performed for each cancer type and independently with mutation status of BRCA1 and BRCA2. Then the cancer types that showed a trend from the results with either gene were selected for the combined analysis. The combined survival analysis with CHAMP1 expression and mutation status of BRCA1 and BRCA2 was performed with tumors being considered mutated if they had a mutated status for either BRCA1 or BRCA2. Also, the Cox proportional hazards regression was performed with accounting for the differences between cancer types and between tumor stages.

### Cancer cell lines’ data acquisition and analysis

The association analyses between cyclin E (CCNE1) expression and CHAMP1 dependency and CHAMP1 expression in cancer cell lines were performed using the expression data from the Cancer Cell Line Encyclopedia (CCLE) project (Ghandi et al., 2019) and the dependency data from the Broad Institute Cancer Dependency Map (DepMap; CRISPR DepMap Public 19Q4 dataset) (Meyers et al., 2017). Both datasets were downloaded from the DepMap Portal (https://depmap.org/portal/). The RNASeq expression counts were normalized by the TMM (weighted trimmed mean of M-values) method using the edgeR package (Robinson et al., 2010) and transformed into log2-counts per million (log2-CPM) values. For each cancer lineage, the low- and high-cyclin E mRNA expressing groups were determined using the median log2-CPM. The significance of the difference in the CHAMP1 dependency between the low- and high-cyclin E mRNA groups were assessed by the Wilcoxon rank sum test using the ggpubr R package. The correlation between CHAMP1 mRNA expression and cyclin E mRNA expression were performed by the simple linear regression on the log2-CPM values using the ggpmisc R package. The plots were generated using the ggplot2 package in R.

