## Supplementary material for "REV7/FANCV Binds to CHAMP1 and Promotes Homologous Recombination Repair": Supplemetal Figures

### Figure S1, related to Fig. 1.

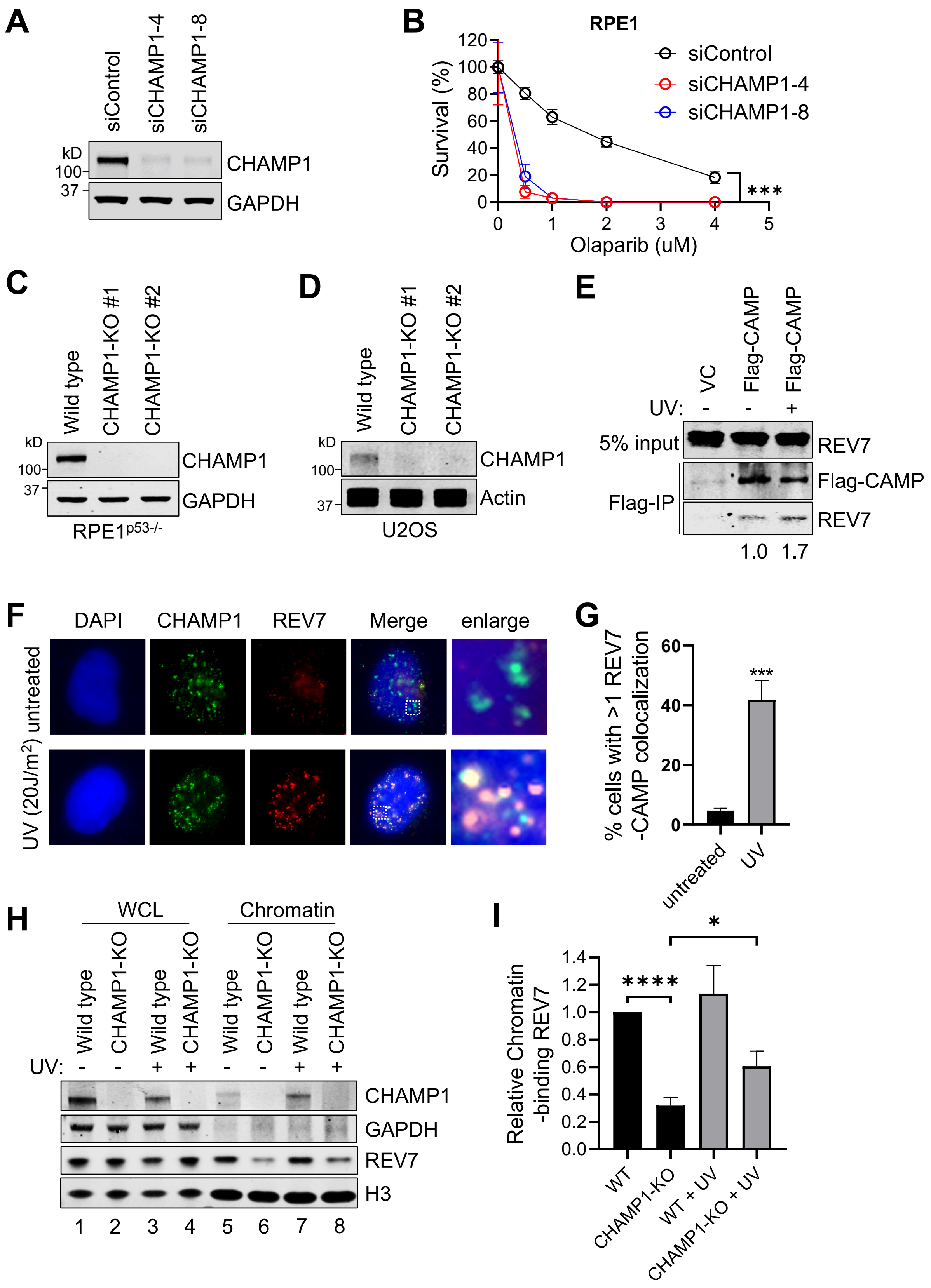

**Supplemental figure 1, related to Figure 1.**

**A**, Western blot showing knocking down efficiency of siRNA targeting CHAMP1. U2OS cells were treated with siRNA negative control or siCHAMP1 for 48hrs. **B**, A 14 days clonogenic assay of RPE1 cells treated with siRNA control and siCHAMP1, and treated with various doses of Olaparib; n=3 independent experiments, ***P<0.0001. Statistical analysis was performed using two-way ANOVA. **C**, Western blot showing the lack of CHAMP1 expression in two CHAMP1 knockout RPE1 cell lines. **D**, Western blot showing the lack of CHAMP1 expression in two CHAMP1 knockout U2OS cell lines. **E**, Western blot showing FLAG-immunoprecipitation of FLAG-CAMP and the co-immunoprecipitation of endogenous REV7 in HEK293T cells with or without UV (20J/$m^2$) treatment. **F**, Representative images of CHAMP1 and REV7 foci 1h after UV (20J/$m^2$) treatment in U2OS cells. co-localizations of CHAMP1 and REV7 are shown as indicated. DAPI was used to stain the nuclei. **G**, Quantification of CHAMP1-REV7 foci co-localization shown in (A). n=3 independent experiments. Error bars indicate standard errors, and p values were calculated using two-tailed Student t-test, ***P<0.0001. **H**, Chromatin fraction of CHAMP1 and REV7 in U2OS wild-type and CHAMP1-KO cells with or without UV (20J/$m^2$) treatment as indicate. Histone H3 is used as control for chromatin isolation. **I**, Quantification of Chromatin-bound REV7 in (G).

**Figure S2, related to Fig. 2.**

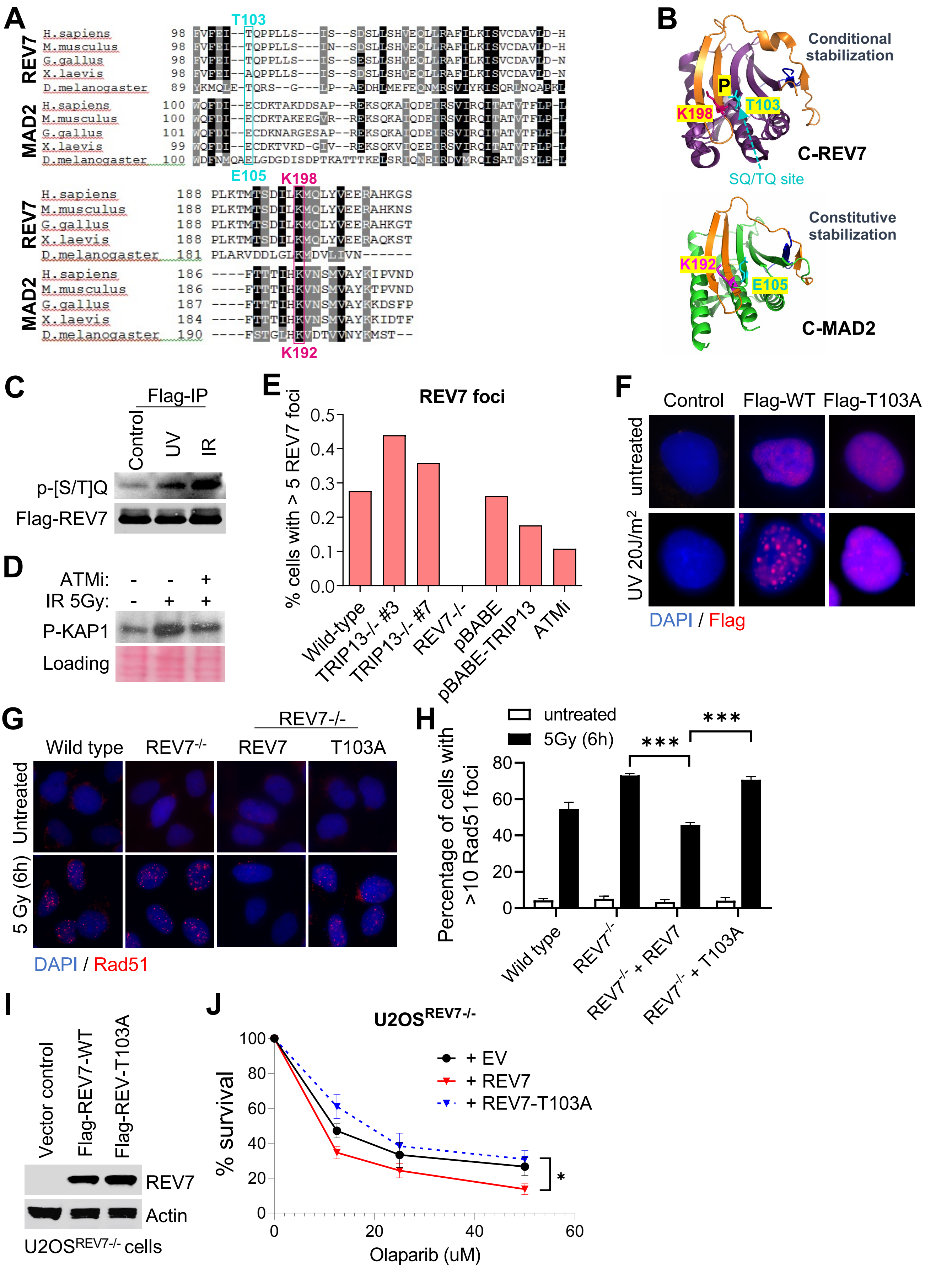

**Supplemental figure 2, related to Figure 2.**

**A**, Primary sequence alignment of REV7 and MAD2 from various organisms. **B**, Structure of the closed form of REV7 and closed form of MAD2. C-REV7 is shown in purple and orange (seatbelt domain). C-MAD2 is shown in green and orange (seatbelt domain). **C**, 293T cells were transfected with FLAG-REV7, and following treatment with UV (20J/$m^2$) for 1 hour or IR (5Gy) for 2 hours. The FLAG-immunoprecipitations were detected by western blot using anti-Flag and anti-p-[S/T]Q antibodies. **D**, Western blot showing phosphor-KAP1 in U2OS cells treated with/without ATM inhibitor, following IR treatment as indicated. **E**, Quantitative analysis of REV7 foci formation in wild type, TRIP13-/-, REV7-/-, TRIP13-overexpressed and ATMi treated U2OS cells after IR (5Gy) treatment. **F**, Representative images of REV7 foci formation in 293T cells expressing Flag-tagged wild type REV7 and REV7-T103A mutant. DAPI was used to stain the nuclei. **G**, REV7-KO U2OS cells were transfected with GFP-Empty Vector, Flag-REV7 wild-type or Flag-T103A mutant, following with/without IR treatment as indicated. **H**, Quantification of >10 RAD51 foci. Error bars indicate standard errors, and p values were calculated using two-tailed Student t-test, ***P<0.0001. **I**, U2OS-REV7$^{-/-}$ cells were transfected with Empty Vector, Flag-REV7 wild-type or REV7-T103A mutant, then detected by western blot using anti-REV7. Actin acts as loading control. **J**. A 14 days clonogenic assay of same cell lines in (I) treated with various doses of Olaparib; n=3 independent experiments, *P<0.05. Statistical analysis was performed using two-way ANOVA.

**Figure S3, related to Fig. 3.**

A

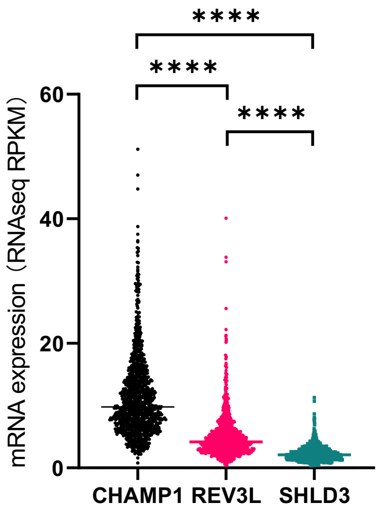


B

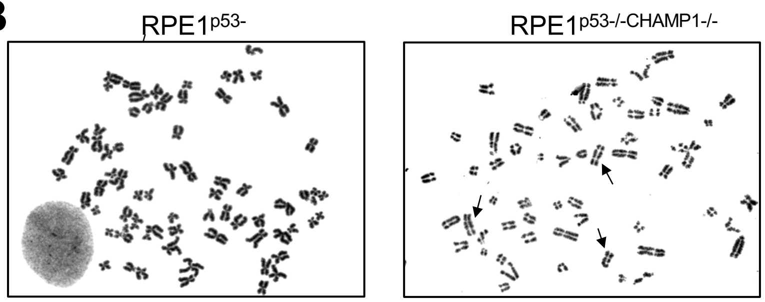


C

| | Chromosome Radials | Chromosome Dicentrics |
|---|---|---|
| RPE1p53-/- | - | - |
| RPE1p53-/-REV7-/- | ++ | - |
| RPE1p53-/-CHAMP1-/- #1 | - | ++ |
| RPE1p53-/-CHAMP1-/- #2 | - | ++ |

D

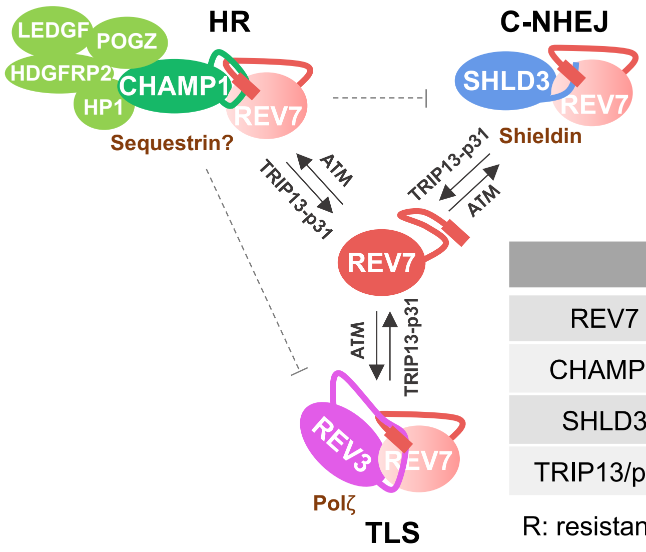


| | Olaparib | MMC |
|---|---|---|
| REV7 KO | R | S |
| CHAMP1 KO | S | R |
| SHLD3 KO | R | R |
| TRIP13/p31 OE | R | S |

R: resistant; S: sensitive

**Supplemental figure 3, related to Figure 3.**

**A,** Relative mRNA expression of CHAMP1, REV3 and SHLD3 genes in all cancer cell lines (Broad, 2019). **B**, Representative images of metaphase spreads from RPE1$^{p53-/-}$ and RPE1$^{p53-/-CHAMP1-/-}$ cells. Arrows show the chromosome dicentrics. **C**, Table summarizing chromosome radial formation and chromosome dicentrics in RPE1$^{p53-/-}$, RPE1$^{p53-/-REV7-/-}$, RPE1$^{p53-/-CHAMP1-/-}$#1, and RPE1$^{p53-/-CHAMP1-/-}$#2 cells. **D**, (left) Schematic of our proposed model of CHAMP1 function in HR, c-NHEJ and TLS regulation. (right) The table showing summary of Olaparib and MMC sensitivity of indicated cells.

**Figure S4, related to Fig. 4.**

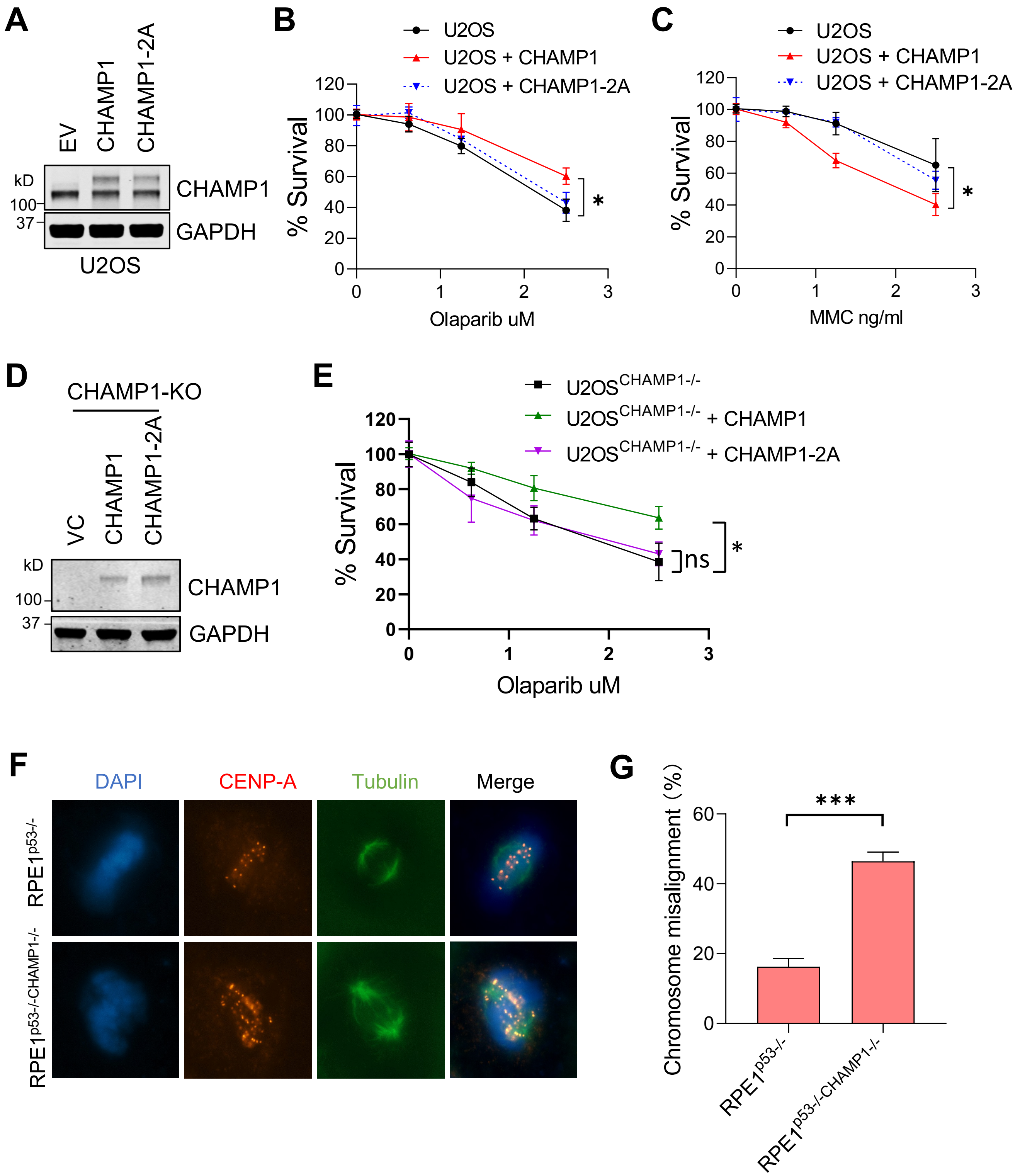

**Supplemental figure 4, related to Figure 4.**

**A**, U2OS cells were transfected with GFP-Empty Vector, CHAMP1 wild-type and CHAMP1-2A mutant. Western blot showing the expression of CHAMP1. GAPDH acts as loading control. A 14 days clonogenic assay of same cell lines in (A) treated with various doses of Olaparib (**B**) or MMC (**C**); n=3 independent experiments, $^*P<0.05$. Statistical analysis was performed using two-way ANOVA. **D**, U2OS$^{CHAMP1-/-}$ cells were transfected with GFP-Empty Vector, GFP-CHAMP1 wild-type or GFP-CHAMP1-2A mutant. Western blot showing the expression of CHAMP1. GAPDH acts as loading control. **E**, A 14 days clonogenic assay of same cell lines in (F) treated with various doses of Olaparib; n=3 independent experiments, $^*P<0.05$. Statistical analysis was performed using two-way ANOVA. **F,** CHAMP1 knockout induced chromosome misalignment. RPE1$^{p53-/-}$ and RPE1$^{p53-/-CHAMP1-/-}$ cells were stained with anti-tubulin and anti-CENP-A. DNA was stained with DAPI. **G**, Quantitative analysis of chromosome misalignment in CHAMP1 depleted RPE1 cells. Error bars indicate standard errors, and p values were calculated using two-tailed Student t-test, $^{***}P<0.0001$.

### Figure S5, related to Fig. 5.

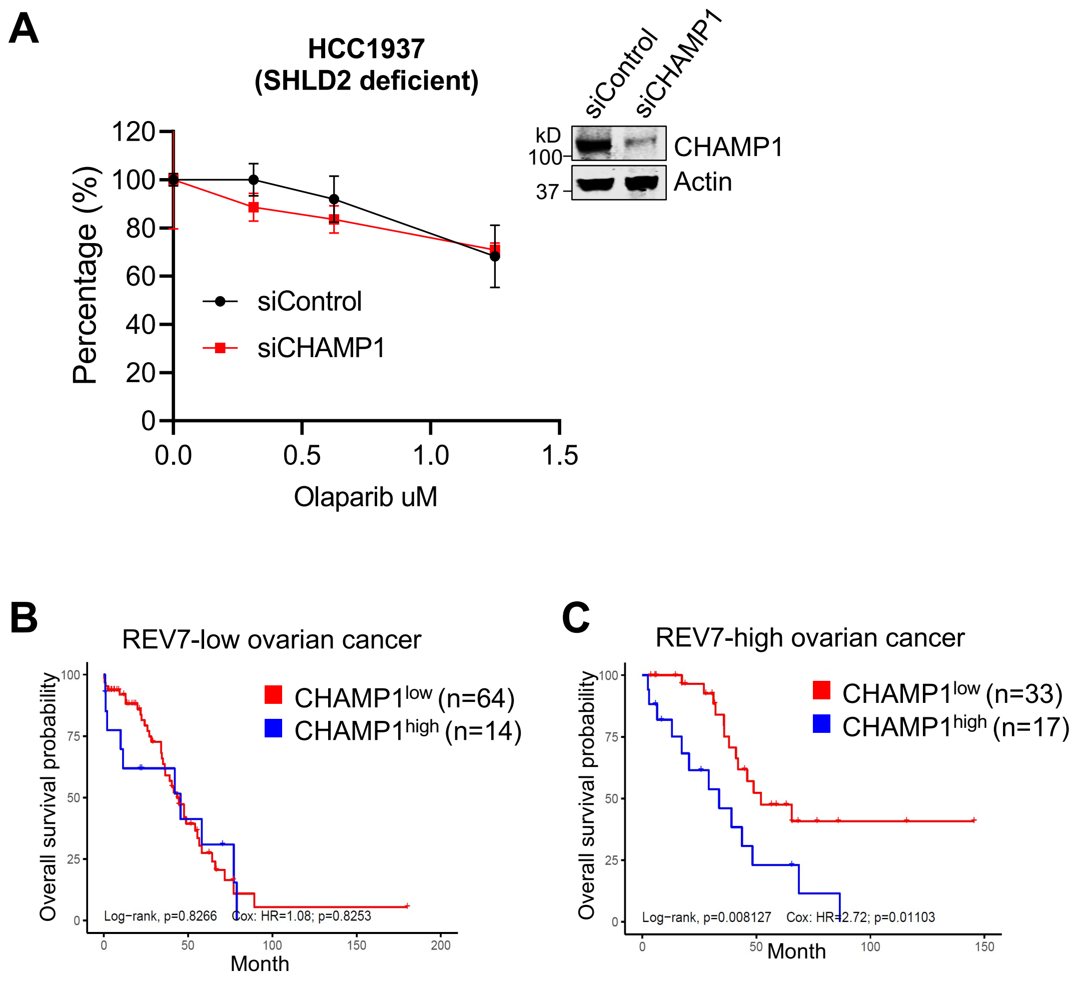


**Supplemental figure 5, related to Figure 5.**

**A**, A 14 days clonogenic assay of HCC1937 cells (SHLD2 deficient) treated with siControl or siCHAMP1 with various doses of Olaparib. Western blot showing the expression of CHAMP1. Actin acts as loading control. **B–C**, Kaplan–Meier curves depicting overall survival of ovarian cancer patients with CHAMP1 expression and REV7 expression.

**Figure S6, related to Fig. 6.**

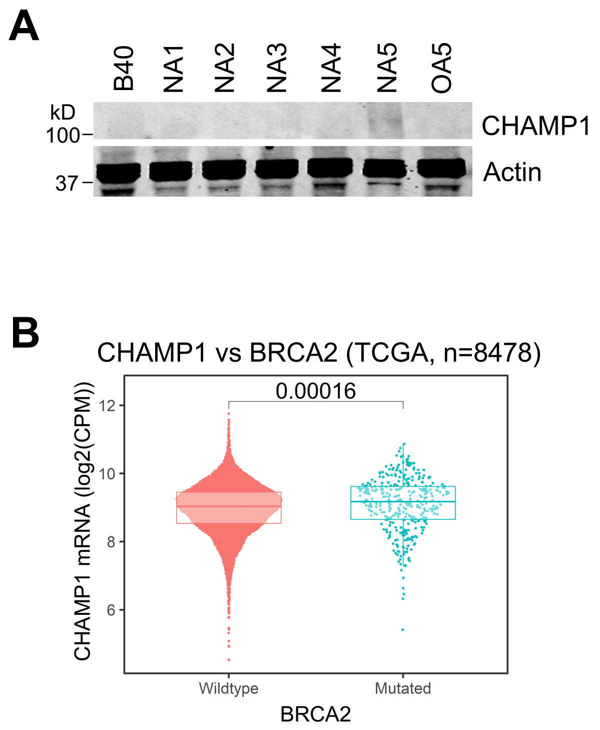


**Supplemental figure 6, related to Figure 6.**

**A**, Western blot showing the expression of CHAMP1 in acquired PRAPi resistant single clones.

**B**, CHAMP1 is highly expressed in BRCA2 mutated tumors.

**Figure S7, related to Fig. 7.**

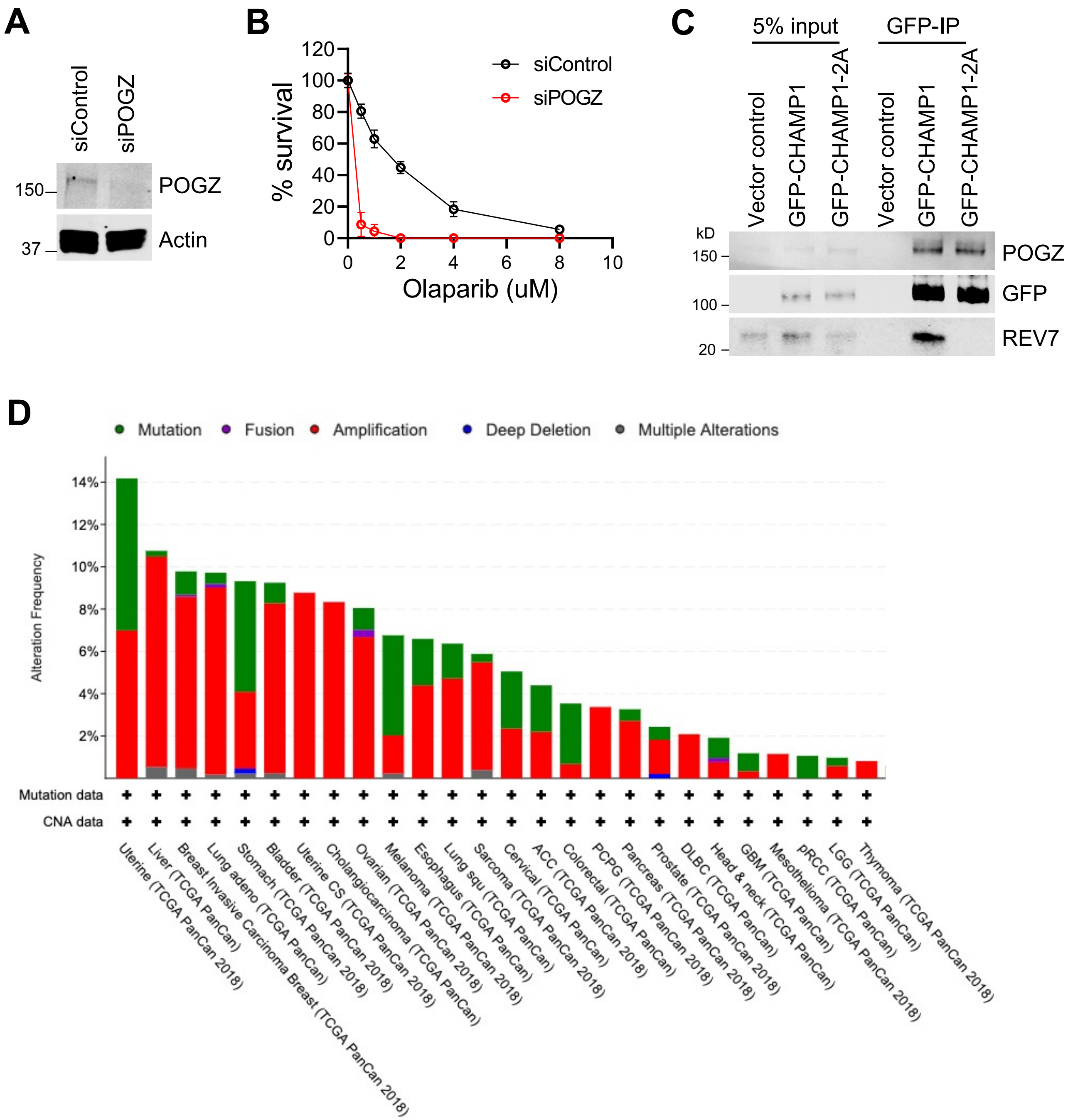

**Supplemental figure 7, related to Figure 7.**

**A**, Western blot showing knocking down efficiency of siRNA targeting POGZ. U2OS cells were treated with siRNA negative control or siCHAMP1 for 48hrs. **B**, A 14 days clonogenic assay of RPE1 cells treated with siRNA control or siCHAMP1 with various doses of Olaparib; n=3 independent experiments, ***P<0.0001. Statistical analysis was performed using two-way ANOVA. **C**, Western blot showing GFP-immunoprecipitation of GFP-CHAMP1 or -CHAMP1-2A mutant, and the co-immunoprecipitation of endogenous CHAMP1 and POGZ in HEK293T cells. **D**, Bar chart showing the prevalence of amplifications (red), deletions (blue), and mutations (green) of the POGZ gene across an array of cancer types in TCGA.
